# Compensatory evolution via cryptic genetic variation: Distinct trajectories to phenotypic and fitness recovery

**DOI:** 10.1101/200725

**Authors:** Sudarshan Chari, Christian Marier, Cody Porter, Emmalee Northrop, Alexandra Belinky, Ian Dworkin

**Affiliations:** Program in Ecology, Evolutionary Biology and Behavior; BEACON Center for the Study of Evolution in Action; Department of Integrative Biology Michigan State University; Lewis-Sigler Institute for Integrative Genomics, Princeton University; Department of Biology, McMaster University

**Keywords:** Compensatory mutations, compensatory evolution, compensatory adaptation, artificial selection, experimental evolution, epistasis, pleiotropy, sexual selection, natural selection, genetic background effects

## Abstract

Populations are constantly exposed to deleterious alleles, most of which are purged via natural selection. However, deleterious fitness effects of alleles can also be suppressed by compensatory adaptation. Compensatory mutations can act directly to reduce deleterious effects of an allele. Alternatively, compensation may also occur by altering other aspects of an organisms’ phenotype or performance, without suppressing the phenotypic effects of the deleterious allele. Moreover, the origin of allelic variation contributing to compensatory adaptation remains poorly understood. Compensatory evolution driven by mutations that arise during the selective process are well studied. However less is known about the role standing (cryptic) genetic variation plays in compensatory adaptation. To address these questions, we examined evolutionary trajectories of natural populations of *Drosophila melanogaster* fixed for mutations that disrupt wing morphology, resulting in deleterious effects on several components of fitness. Lineages subjected only to natural selection, evolved modifications to courtship behavior and several life history traits without compensation in wing morphology. Yet, we observed rapid phenotypic compensation of wing morphology under artificial selection, consistent with segregating variation for compensatory alleles. We show that alleles contributing to compensation of wing morphology have deleterious effects on other fitness components. These results demonstrate the potential for multiple independent avenues for rapid compensatory adaptation from standing genetic variation, which ultimately may reveal novel adaptive trajectories.

## Introduction

Populations are constantly exposed to deleterious mutations (1–3). It is likely that most deleterious alleles are kept at low frequency or purged from populations by natural selection. However deleterious mutations can rise in frequency and even fix in populations due to genetic drift, hitchhiking with beneficial alleles or antagonistic pleiotropy, among other mechanisms (4–7). In such cases, the deleterious consequences of such a mutation can be suppressed via epistatic compensatory mutations. Compensatory mutations conditionally reduce the fitness cost of the original deleterious mutation, which is well documented in microbial (4, 8–13) and a few animal systems (5, 14–16).

Mutations are often pleiotropic, and it is unclear whether compensatory adaptation proceeds by directly or indirectly ameliorating one or more phenotypic aspects of the deleterious mutation. For instance, the potential decline in male courtship song due to an allele reducing wing stridulation (due to predator induced selection) in crickets was compensated by evolving a modified mating strategy (17–19). Given that traits are often utilized for multiple aspects of organismal performance, it is unclear whether compensation for one of the functions will recover others or whether epistatic pleiotropy (20, 21) will restrict the compensatory response.

Experimental studies of compensatory evolution are often initiated with a single (low fitness) genotype. Experimental evolution proceeds, recovering fitness via *de novo* mutations that occur concurrent with natural selection (4, 8, 14). But natural populations may harbour standing genetic variation that can modify phenotypic consequences of mutant alleles and thus potentially contribute to compensatory adaptation. In *Drosophila melanogaster*, artificial selection has been demonstrated to rapidly enhance or suppress specific phenotypic effects of mutations (22–24). Yet, given the known pleiotropic nature of many deleterious alleles, it is unclear whether this would result in complete suppression of the deleterious effects. Alternatively, compensation for the reduction in performance and fitness may occur by evolution in traits that the initial deleterious allele did not directly influence (11, 25). i.e. evolution proceeds by altering other phenotypes that bypass aspects of function for the initially affected one.

We thus wanted to understand how compensation for fitness loss occurred given a complex suite of phenotypes. Would compensatory evolution be the result of direct suppression of the deleterious effects of the focal allele? Or, alternatively, would evolution occur in other traits that compensate for specific fitness components, not initially influenced by the deleterious mutation. Finally, we wanted to determine how segregating (cryptic) variation would be utilized during compensatory evolution. To address these questions we fixed individual mutations in several genes that disrupt aspects of normal wing development in *Drosophila*. The *Drosophila* wing is essential for flight performance, provides both visual and auditory signals (song) during courtship and aggression (26–29) with wing morphology likely being sexually selected. Despite the long-term stability of the venation pattern and overall shape of the *Drosophila* wing (30–32), alleles that qualitatively perturb wing morphology are commonly observed at low frequency in natural populations of *D. melanogaster*. Furthermore, recent work has demonstrated that populations that have rapidly adapted to high altitudes are highly sensitive to mutational perturbation and show an elevated frequency (~30%) of wing abnormalities, potentially due to hitchhiking or the pleiotropic consequence of strong directional selection for other traits (likely increased size) (33). With this in mind, we utilized several mutations. The *vestigial* allele, (*vg^1^*) results in a truncated wing while mutations in the *rhomboid* (*rho^ve-1^*) and *net* (*net^1^*) genes result in wing venation defects as observed at low frequency in natural (Fig. 1A) populations (33). Using the mutant populations, we created replicated treatments of artificial selection, selecting only for phenotypic compensation of the wing (i.e. recovery of the wild-type phenotype). We also generated replicated experimental evolution treatments, with natural (and sexual), but no artificial selection operating. In the artificial selection lineages, we observed rapid and almost complete compensation of wild type wing morphology, consistent with segregating variation for compensatory alleles. Interestingly, there was no wing recovery in the experimental evolution lineages. In these lineages, we observed rapid evolution in aspects of mating behaviour as well as other compensatory response for other fitness components. Thus we demonstrate that fitness recovery can occur via both direct and indirect effects on the phenotypes influenced by the deleterious mutation and there is considerable standing genetic variation for both phenotypic and fitness compensation.

**Fig. 1.**
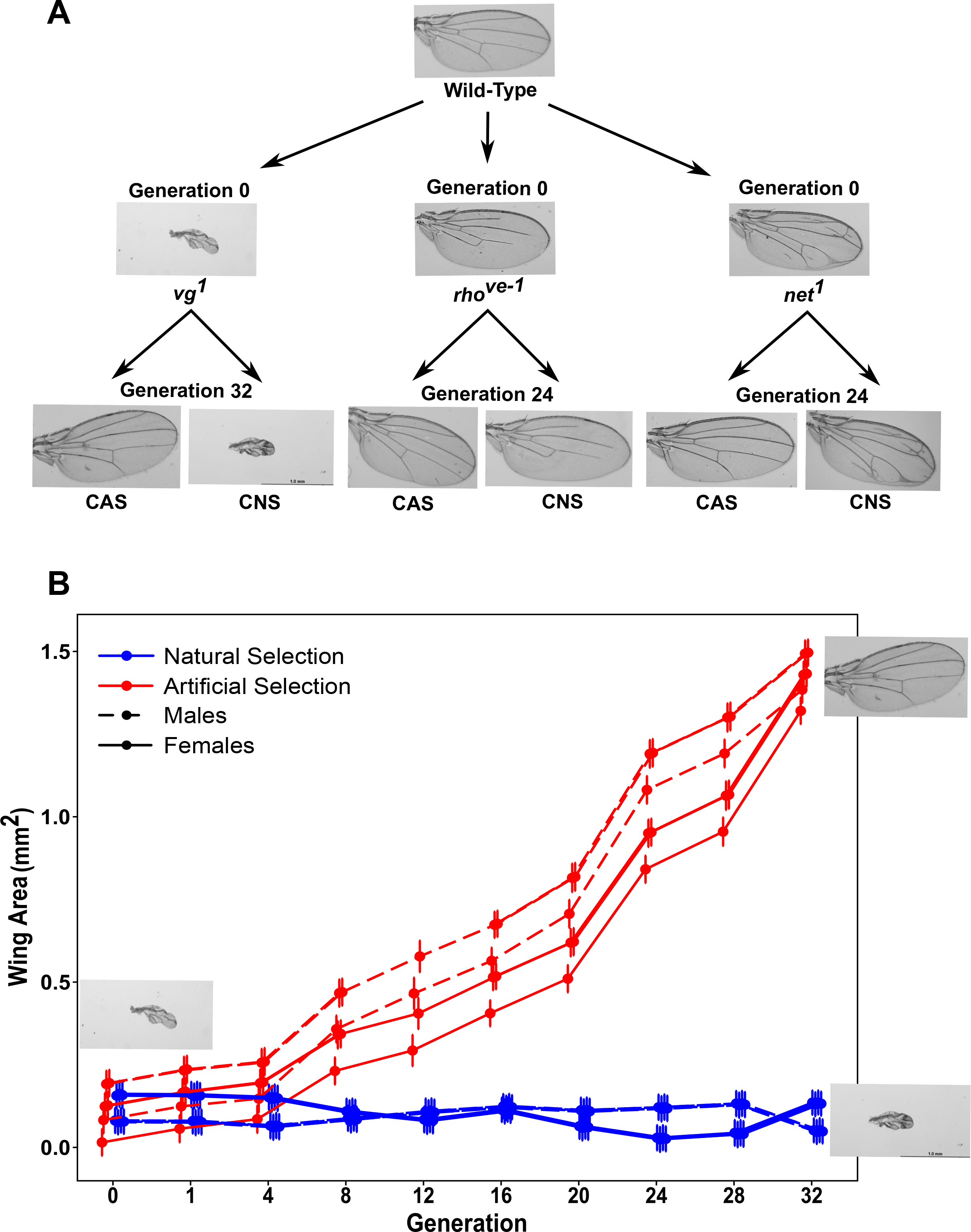
Evolutionary response of wing morphology under different selection regimes. (A) Qualitative images of wings of the wild-type flies, mutants at generation 0 and at later generation under artificial selection (CAS) showing morphological compensation and experimental evolution (CNS), showing no compensation (B) Quantitative evolutionary response of wing morphology in 3 replicates of *vg*^*1*^ CAS and 4 replicates of *vg*^*1*^ CNS populations over 32 generations. The data points and error bars represent the mean wing area and 95% CI of 12-15 wings (individuals)/ treatment/sex/replicate.

## Materials and methods

### Drosophila strains

#### Mutations

The autosomal mutations *vestigial*^*1*^ (*vg*^1^), rhomboid^ve-1^ (*rho^ve-1^)*, and *net^1^* (Fig. 1A and Table S1) were obtained from the Bloomington stock center. *vg*^*1*^ was introgressed into a synthetic outbred population prior to being introgressed into the natural population described below.

#### Maintenance of the *Drosophila* Populations

Adults from all the populations described below were maintained either in large (32.5cm^3^, BugDorm BD43030F) or small (17 cm^3^, BugDorm BD41415) cages at 23°C (+/− 1°C), 30-50% Relative Humidity (RH) and 12hr Light/Dark (L/D) cycle. They were allowed to lay eggs in bottles with 50-60ml molasses-cornmeal media. After egg laying, the bottles were removed from the cages and maintained in a Percival (Model: I41VLC8) incubator at 24°C, 65% RH and 12hr L/D cycle throughout larval stages. For certain populations (described below), the adults upon eclosion and subsequent selection were maintained in a Percival (Model: I41VLC8) incubator at 18°C with 65% RH and 12hr Light/Dark cycle for 5-7 days before being introduced into cages. Fly collections for selection and other assays were performed under light CO_2_ anaesthesia on a Leica M125 microscope.

#### Progenitor *Drosophila* Population

Our population was initiated using more than 500 single pair matings of wild caught *D. melanogaster* individuals from Fennville Winery, MI (GPS co-ordinates: 42.578919, −86.144936). We screened out and discarded *D. simulans* (~5% frequency), and introduced ~1000 progeny from the single pairs (2 males and 2 females from each mating) into a large cage to establish the FVW Ancestral (FVWA) population. Adults were allowed to lay eggs in 10 bottles for 2-3 days and the bottles were transferred to the 24°C incubator. After adults began to emerge, bottles were transferred into a fresh cage and the eclosion process was allowed to occur for 10-12 additional days. After eclosion, old bottles were discarded and population sized reduced to ~1500-3000 individuals, that formed the breeders for the next generation.

#### Introgression of mutant alleles into the FVWA population

We wanted to include alleles that were potentially rare in natural populations (and thus could be rapidly lost during lab domestication) as they may be important for any compensatory response. Thus after 2 generations of the establishment of the FVW population of *D. melanogaster* in the lab we began backcrossing the mutations. Introgression of the mutations into FVW was performed by repeated backcrossing to form a ‘BASE’ mutant population per mutation. Each backcross cycle consisted of mating ~150-200 mutant males to 300-400 virgin FVW females to create F1 hybrids. 600-800 randomly chosen F1 flies were mated to recover recombinant F2 mutant homozygotes. We performed 10 cycles of the backcross for *vg*^*1*^, and 8 cycles each for *rho*^*ve-1*^ and *net*^*1*^. This generates populations of flies that should be segregating much of the natural variation in the FVW population except for near the focal locus itself. Base mutant populations were maintained in small cages at an adult population size of 300-400 flies to minimize natural selection (lab domestication). From the Base mutant population (for each of the mutations), 4 replicates of <u>N</u>atural <u>S</u>election (C<u>NS</u>) lineage, 3 replicates of <u>A</u>rtificial <u>S</u>election (C<u>AS</u>) lineage and 3 replicates of population-size matched <u>C</u>ontrol for <u>A</u>rtificial <u>S</u>election (N<u>ASC</u>) lineage were generated. In addition, 3 replicates of FVW controls were also created to control for natural selection to the overall lab rearing conditions. All adult flies were maintained in small cages in similar conditions as FVWA.

### Selection Procedure

#### Natural Selection (CNS) & FVW Controls

Lineages were initiated by allowing 500 adults to lay eggs for 5-7 days in 4 bottles and allowed to develop in the 24°C incubator. 2-4 days after eclosion of the first adult flies, bottles were transferred to a 17cm^3^ cage for 5-7 days, and then bottles were replaced with fresh, yeasted bottles for egg laying. For CNS lineages, no direct control of density was performed and the populations had very high adult and larval densities. The average population size for *vg*^*1*^= ~2000, *rho*^*ve-1*^ & *net*^*1*^ = ~3000 based on census every 8-10 generations (see Supplementary materials for details). For the FVW controls, the population size was estimated to be ~4000. After females laid eggs, and larvae emerged (to form the next generation), adults were stored in 70% ethanol. We re-introgressed the base mutant stock to the original lab adapted, natural, wild-type population (FVWA) every 8-10 generations for 1-2 backcross cycles as described above, in particular before any comparative testing assays to generate the equivalent of an ‘unevolved ancestral control’ BASE mutant population. For each mutation, we had 4 independently experimentally evolved replicate populations. It is worth noting that with our experimental set-up (in cages), natural selection for flight is likely weak relative to other selective forces (such as sexual selection).

#### Artificial selection (CAS) & Non-selection Controls (NASC)

Adult flies were allowed to mate and lay eggs for 2 days in 5-6 bottles following which, development occurred in the 24°C incubator. We screened ~1000-1200 individuals and selected those that had the least severe manifestation of the mutant wing phenotype. At each generation, *vg*^*1*^ CAS flies were selected for longest and widest wings, while *rho*^*ve-1*^ and *net*^*1*^ were selected for the most (relative to other individuals from that generation) “wild type” venation patterns. From this, 55 pairs of the selected individuals were used for breeding. The NASC controls in this case, were formed by allowing ~55 pairs of adults that were randomly chosen out of ~1000-1200 to breed for the next generation. In both cases the collected flies were maintained at 18°C for 5-7 days before being introduced into cages. After every round of egg laying the adults were stored in 70%. For each mutation, we had three independent evolving artificial selection lineages. During the selection process, genetic tests were performed to confirm that the focal allele (*vg*^*1*^, *rho*^*ve-1*^, *net*^*1*^) remained fixed for each of the relevant populations.

#### Measuring wing and body (thorax) size

To document changes in wing size through the evolutionary process, a single wing was dissected from each of 15 individuals/sex/population/selection regime/replicate and mounted in 70% glycerol/30% PBS (with phenol as a preservative) for a total of 30 observations/ replicate. For the CNS and CAS populations with *vg*^*1*^, the dissection was performed every 4 generation up to generation 32, while for *rho*^*ve-1*^ and *net*^*1*^ it was done every 8 generations, up to 24. For the BASE, FVW and NASC lineages, the dissections were performed at generation 0 and either generation 32 or 24 respectively. To document changes in wing size for the *vg*^*1*^ within a generation at low and high densities, a single wing was dissected from a minimum of 10 and a maximum of 20 individuals/sex/population/replicate/density/vial at generation 27. Images of the wings were captured using an Olympus DP30BW camera mounted on an Olympus BW51 microscope using DP controller image capture software (v3.1.1). The wing area was then obtained using a custom macro in ImageJ software (v1.43u). We imaged the thorax of every fly prior to dissection and measured the body size as the length of the thorax using ImageJ software (v1.43u).

### All the following experiments were performed using vg^1^ and FVW populations

#### Estimation of strength of selection on *vg*^*1*^

We performed an assay to estimate the selection coefficient for the *vg*^*1*^ mutation in the BASE population under conditions with and without mate choice. Each treatment was initiated with three replicates of 100 males and 100 females with 98 being *vg^1^/ vg^1^*, 18 wt/wt and 84 *vg*^*1*^/ wt (initial frequency of *vg*^*1*^ = 0.7). We used the ancestral FVWA population for the wild-type *vg*^*+*^ alleles. For each replicate in the mate choice treatment, we used 20 vials, each with 5 males and 5 virgin females. For each replicate in the no mate choice treatment we used 100 vials with single mating pairs. In both cases, flies were randomly assigned to a vial. The flies were allowed to mate for 3 days at 24°C, following which the males were discarded and the females were transferred to small cages. They were allowed to oviposit for 4 days in 4 lightly yeasted bottles. Females were discarded and the bottles placed at 24°C. Upon eclosion, males and virgin females were collected for a period of 4 days and scored. After collection and census, progeny were randomly placed into vials with or without mate choice as described above. Assuming Hardy-Weinberg equilibrium we estimated the *vg*^*1*^ allele frequency from the frequency of *vg*^*1*^ in the census. In addition, to independently estimate *vg*^*1*^ allele frequency, we performed a test cross every 3 generations by independently mating 50 phenotypically wild-type females from each replicate to homozygous *vg*^*1*^ male. We calculated the selection coefficient per generation [s = 1-(q’/q)] and averaged throughout the duration of the experiment. To analyse the rate of loss of the mutant allele in different populations over multiple generations, we fit a linear model with fixed effects of population, generation and their interaction.

#### Estimation of selection on the *vg*^*1*^ Wing

Previous evidence has demonstrated that wing size, shape and interference patterns are target of selections with respect to female mate choice (26–29). To determine the extent to which phenotypic variation in wing size for *vg*^*1*^ individuals remained a target of selection, we first generated an F1 panel by reciprocally mating equal numbers of males or females CAS flies from replicate 2 at generation 33 to the BASE population in bottles for 3-4 days. The resulting F1 population encompassed the entire variation spectrum from being phenotypically *vg*^*1*^ like (5-10% wild type wing area) to having almost wild-type like wings (100%). We then qualitatively screened for equal numbers virgin females and males that encompassed the entire wing size variation spectrum and introduced them into a large cage to breed for the F2 generation for 3-4 days. The resulting F2 population also encompassed the entire wing size variation spectrum. Using flies from the F2 population, we performed 204 choice assays by providing a female from the base population with 2 males- one with a 5-10% (of wild type size) wing and another with a 10-15% wing per assay. All assays were performed in vials with 10ml food. Virgin males and females were collected and 3-6 days post eclosion, age matched flies were randomly aspirated into vials for the assay. Successful mating pairs were separated and the wing area of the left and right wing as well as body size of both the successful and unsuccessful males were measured. This was fit as a generalized linear mixed model (logistic) with fixed effects of male and female wing areas (and their interaction) and the intercept was allowed to vary according to the random effect of trial nested within block.

#### Behavioural and Life-history assays

Prior to all assays described below, the base populations were reintrogressed for at least one generation (irrespective of previous introgressions) into the FVWA population. Also, ~1000 flies from each population were collected separately and maintained under common conditions for one generation to eliminate parental effects. : Eggs for all of life-history assays were collected on grape juice agar plates with 50-60% grape juice and 2% agar.

#### Mating Assays

We performed courtship assays for the artificial selection, natural selection (fixed with *vg*^1^), BASE *vg*^*1*^ and FVW control populations. Each of the populations was reared at moderate-high larval rearing density (to simulate conditions during experimental evolution). Once adults emerged, they were communally maintained as 15 virgin males or virgin females per vial). They were age matched (3-6 days post-eclosion) and received identical conditions for light, food and humidity. For the assay we introduced 15 pairs into a 17cm^3^ cage with a small amount of food and scored the number of flies courting or copulating every 5 minutes for a total duration of 70 minutes. In order to differentiate between male lineage or treatment specific patterns of male persistence choice (or preferences), we performed similar cage courtship assays between a specific sex from the selection lineages and the corresponding opposite sex from each other treatment lineage. We modelled the maximum proportion of males courting (i.e. across all time points for a given cage) using a linear mixed model with the fixed effects of both female and male source populations and a random effect for the date of the assay. In this case replicate effects was negligible and hence not included in the final model. We modeled the maximum proportion of males copulating for a given cage using a linear mixed model with the fixed effects of both female and male source populations. In this case we also fit three independent random effects of the replicate population nested within female, within male and the effect of date of the assay.

#### Larval Competitive ability

Eggs from every replicate of FVW, CAS, CNS, NASC and Base populations were placed in a food vial (~10ml) with equal number of eggs from a common competitor population (Inbred, lab strain: Samarkand wild-type marked with *white^-^* allele) at low (25+25 eggs) and high (150+150 eggs) densities. Upon emergence, we scored flies for red or white (common competitor) eye-colour. We also performed a similar experiment, competing CNS and BASE populations against FVW populations as a common competitor (scored based on wing morphology). To analyse the survivorship of the treatment flies over the common competitor, we fit a logistic regression with fixed effects of density and population, with replicate nested within population, and the interaction between population and density. We also fit fixed effects for vial nested within block.

#### Egg to adult viability and development time

Eggs from every replicate of FVW, CAS, CNS, NASC and Base populations were placed in a food vial at low (50 eggs) and high (300 eggs) densities. To determine viability, we scored and calculated the proportion of flies that eclosed. We also recorded development time (days to eclosion). To analyse egg to adult survivorship, we fit a linear model with fixed effects of density and population, with replicate nested within population, and the interaction between population and density. We also fit fixed effects for vial nested within block. To analyse the egg to adult development time, we fit a linear model with fixed effects of population with replicate nested within population, density and sex and all the two-way interaction between these. We additionally fit fixed effects of block and vial nested within year.

#### Female Fecundity

20 virgin females from every replicate of FVW, CAS, CNS, NASC and Base populations were introduced with conspecific virgin males 2-3 days after eclosion, as single pairs in a lightly yeasted vial (<10ml food) for 3 days. This three-day mating period was to mimic (in part) the waiting time in selection treatments before new bottles are added. After this 3-day mating period, each single pair was transferred to a fresh vial (~10ml food) to lay eggs- termed as day 1− for 24 hours. On day 2, the single pair was transferred to another minimally yeasted fresh food vial. This single pair was transferred into fresh food vial without yeast for another 2 days. To determine female fecundity we counted the total number of eggs laid over 4 days. We fit a linear model with fixed effects of population, with replicate nested within population, density, body size and all the two-way interaction between these. We additionally fit a fixed effect for block.

#### Statistical analyses

All statistical analysis were performed in R v3.1.1 using lm(), glm() (base R) and lmer() (lme4 library v1.1-12). This was followed by extracting and plotting the relevant coefficients using the effects package v3.0-0. Other plots were generated using the ggplot2 package v2.1.0.

## Results

### Rapid compensation to near wild type wing morphology under artificial selection is consistent with segregating variation for suppressor mutations

Despite the severe consequences of the mutational perturbation on wing morphology in the ancestral (base) population, we observed rapid recovery of “normal” wing morphology in lineages artificially selected for “wild type” like wing morphologies. This included compensation of wing morphology for the almost complete loss of the wing blade observed in ancestral *vg*^*1*^ population (Fig. 1A and 1B). While the response to selection was continuous, near phenotypically wild-type wings were observed at low frequency by the 14^th^ generation for the lineages fixed with the *vg*^*1*^ mutation. By generation 32, mean wing area increased to ~1.43 mm^2^ (CI 1.39, 1.46) from the initial ~0.12 mm^2^ (CI 0.089, 0.16). This compensatory response resulted in wings only slightly smaller than what we observed for our wild-type sizes of ~1.5mm^2^ (CI 1.47, 1.54), averaged across sexes. The increase in wing size was not correlated with increases in body size, which remained relatively consistent throughout the evolutionary process (Fig. S1A, B). Interestingly, halteres (which are also reduced in vg^1^) demonstrated a correlated compensatory response to artificial selection (not shown). Similar rapid wing-phenotype recovery in *rho*^*ve-1*^ and *net*^*1*^ CAS lineages was also observed (Fig. 1A).

### Experimental evolution under natural selection did not result in the recovery of wild type wing morphology

Despite segregating variation for compensation of wild type like wing morphology (under artificial selection), we did not observe any recovery of the wings in the experimentally evolved CNS lineages (Fig. 1A and 1B). Similarly, populations allowed to evolve by natural selection fixed for the *rho* and *net* alleles also did not demonstrate any phenotypic compensation of the wing venation defects (Fig. 1A).

### *vg*^*1*^ has deleterious effects on multiple fitness components

Given that the natural selection (CNS) lineages did not show phenotypic compensation of wing morphology, we sought to address why no response was observed. Results from the artificial selection treatments demonstrate that there is no genetic constraint on response, and that segregating variation in the population is sufficient to compensate for the phenotypic effects of the mutations on wing morphology. This suggests that the likely explanation for the lack of phenotypic evolution under natural selection may have been due to weak natural selection, countervailing selective pressures or antagonistic pleiotropy. Thus we examined the effects *vg*^*1*^ had on several fitness components. Since wings are involved in sexual signalling (26–29), we determined the extent to which selection on mate choice would influence the deleterious nature of the *vg*^*1*^ allele. We examined the allelic dynamics in populations initiated at high frequency (but not fixed) for the *vg*^*1*^ allele, with and without opportunity for mate choice. The frequency of the *vg*^*1*^ allele rapidly decreased in all treatments indicating sexual selection dependent and independent deleterious effects on fitness (Fig. 2A and Fig. S2A). Furthermore, selection against *vg*^*1*^ was stronger in the presence of mate choice (mean selection coefficient ~0.26 and CI 0.18, 0.34 in presence of mate choice while it was ~0.13 and CI 0.048, 0.33 without mate choice), consistent with the known role of wings in sexual selection. The deleterious nature of the *vg*^*1*^ allele on mate choice may be a direct impact of the reduction in wing size itself (28), and “within population” mate choice experiments (Fig. S2B) were consistent with very weak sexual selection on wing size even within populations fixed for the *vg*^*1*^ allele.

**Fig. 2.**
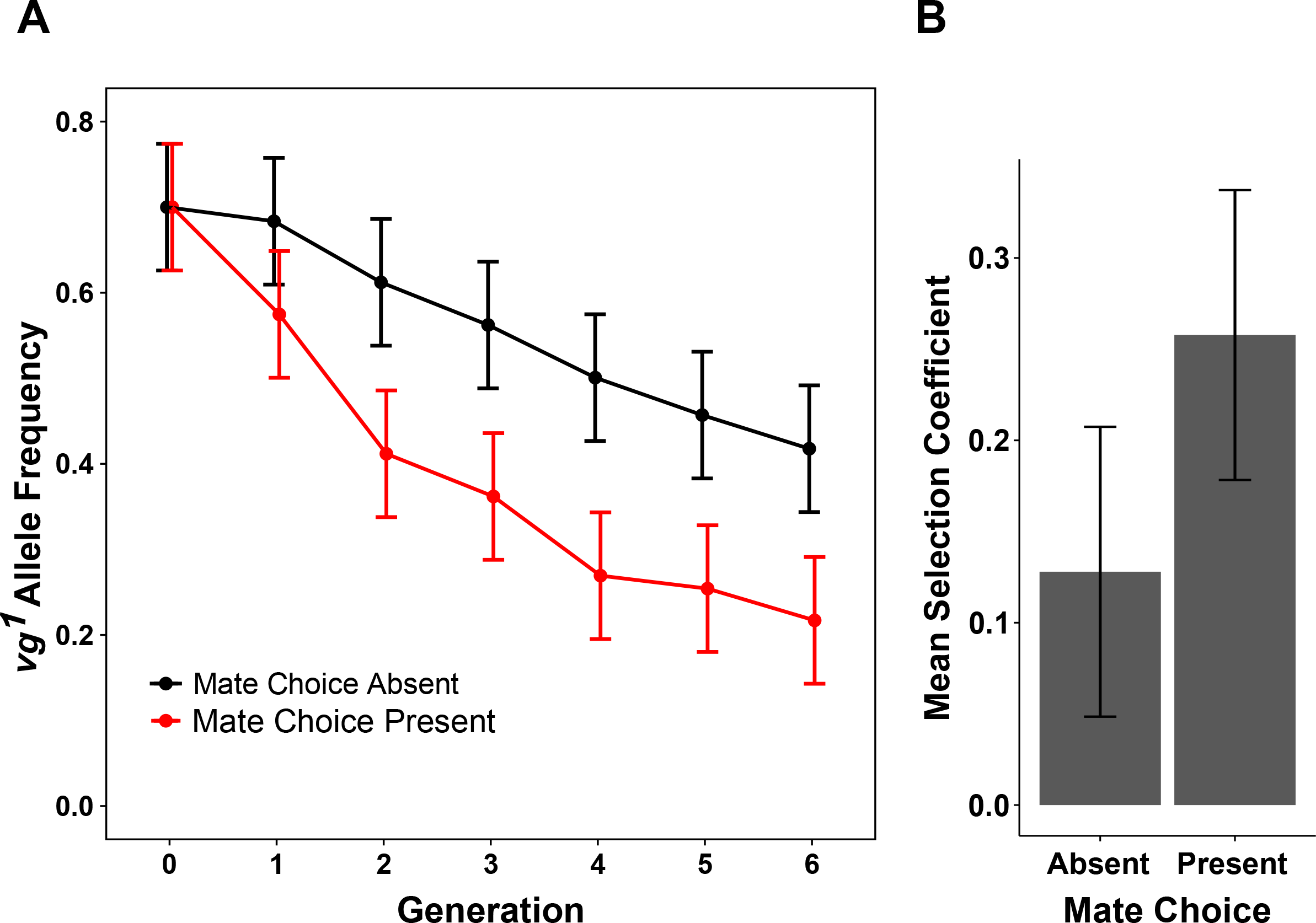
Sexual selection dependent and independent deleterious effects of the *vg*^*1*^ allele. (A) Faster extinction of the *vg*^*1*^ allele in the presence of mate choice as compared to the absence of mate choice. The data points and error bars represent the mean *vg*^*1*^ allele frequency and 95% CI of 3 replicates of each treatment. (B) Stronger selection against *vg*^*1*^ in presence of mate choice as compared to the absence of mate choice. The bars represent mean selection coefficient and error bars represent 95% CI of 3 replicates of each treatment. Each of the replicate was initiated with a population of 200 individuals that subsequently expanded to around 600-900 individuals.

### Behavioural adaptations may compensate for altered courtship signals in the experimentally evolved lineages of *vg*^*1*^

We tested whether the experimentally evolved (CNS) populations, which lacked the ability to generate aspects of the wing-mediated sexual signals, compensated by altering other aspects of mating behaviour. Mating assays demonstrated that a higher proportion of males from the experimentally evolved CNS populations were engaged in courtship at any given time compared with other populations (Fig. 3A, Fig. S3A and Fig. S4). This increase in courtship was associated with increased copulation success in CNS lineages, compared to the ancestral BASE populations although it did not recover “wild type” levels (Fig. 3B, Fig. S3B and Fig. S5). We also observed that the proportion of artificially selected CAS flies courting (Fig. 3A, Fig. S3A and Fig. S4) were similar to the control populations without the *vg*^*1*^ mutation with an intermediate level of copulation (Fig. 3B, Fig. S3B and Fig. S5), consistent with wing morphology being a target of sexual selection.

**Fig. 3.**
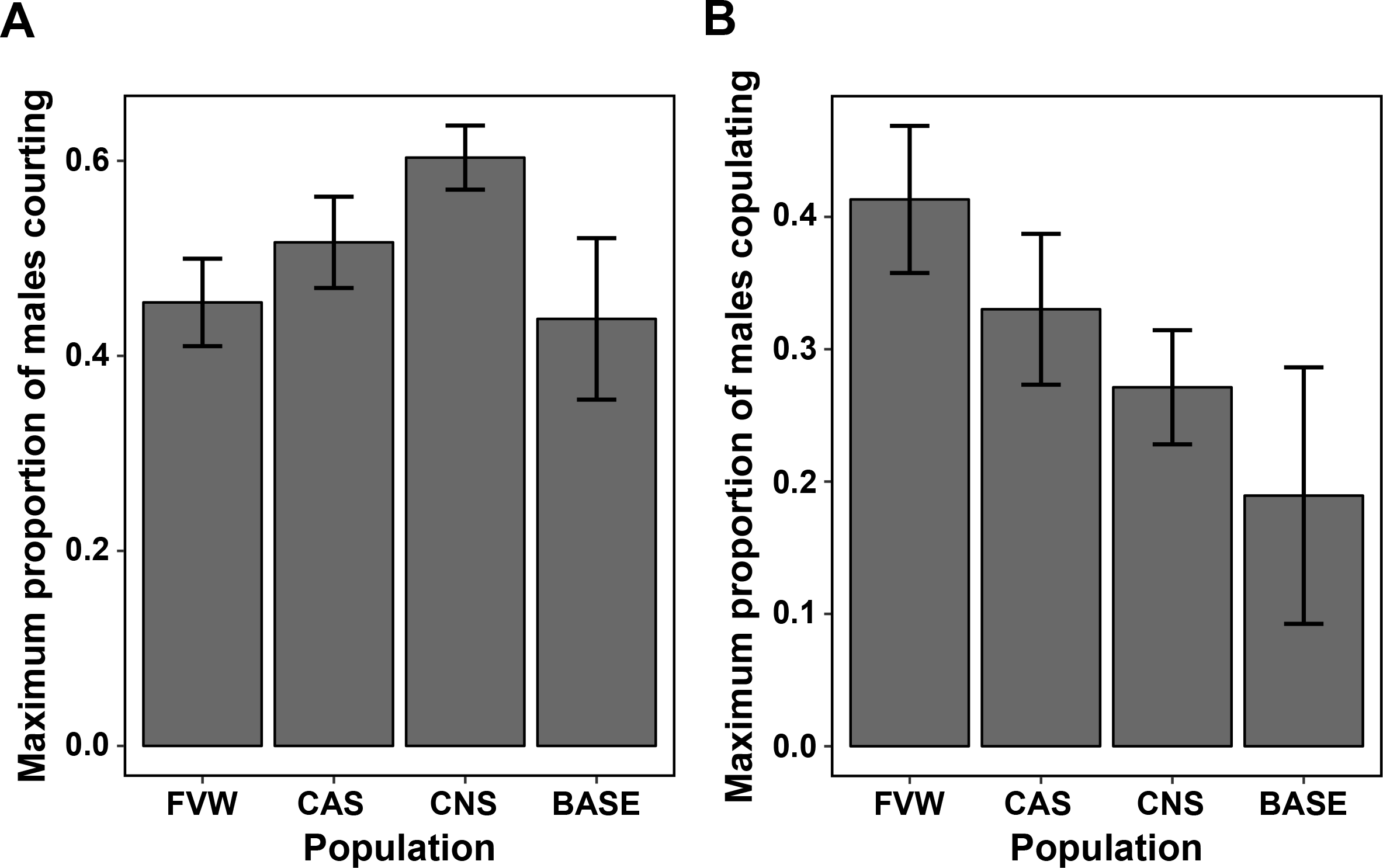
Behavioral compensation in the mating behavior of populations under different selection regimes. (A) Maximum courtship proportion of males and females from each of the populations with CNS population displaying highly persistent courtship behavior (B) Maximum copulation success for each of the populations with CNS population displaying higher mating success than the unevolved BASE population and CAS displaying high mating success mediated via the recovery of the wing phenotype. The data points and error bars represent the mean proportion of males courting or copulating and 95% CI, for 15 male-female pairs of 3 replicates of FVW wild-type, 3 replicates of *vg*^*1*^ CAS, 4 replicates of *vg*^*1*^ CNS and 1 replicate of *vg*^*1*^ BASE populations averaged over 70 minutes.

To assess whether the changes in the mating behaviours in the CNS lineages due to an evolved increase in male persistence, decrease in female choosiness or both, we examined response across all possible combinations of populations. We observed that CNS males consistently courted more than all other lineages, irrespective of the lineage of the females they were courting (Fig. 3A, Fig. S3A and Fig. S4). This indicated that behavioural changes had occurred in part due to increased male persistence in the CNS populations.

### Experimentally evolved lineages compensate for deleterious effects of *vg*^*1*^ on competitive ability and viability, while artificial selection for wing morphology entails a cost

The *vg*^*1*^ allele had a deleterious effect on egg to adult viability in the ancestral base population relative to the FVW controls without the mutant allele (Fig. 4A and Fig. S6A). We observed that egg to adult survivorship increased in the experimentally evolved (CNS) populations compared to the ancestral BASE population. Consistent with a potential cost to the alleles compensating for recovery of wing morphology, egg to adult survivorship was reduced in the artificial selection (CAS) lineages (Fig. 4A and Fig. S6A), compared to almost all populations at both densities, including its population size matched, control lineages for artificial selection (NASC). We observed minimal differences between the wild-type controls and the ancestral wild-type flies at high density while it was more variable at low density.

**Fig. 4.**
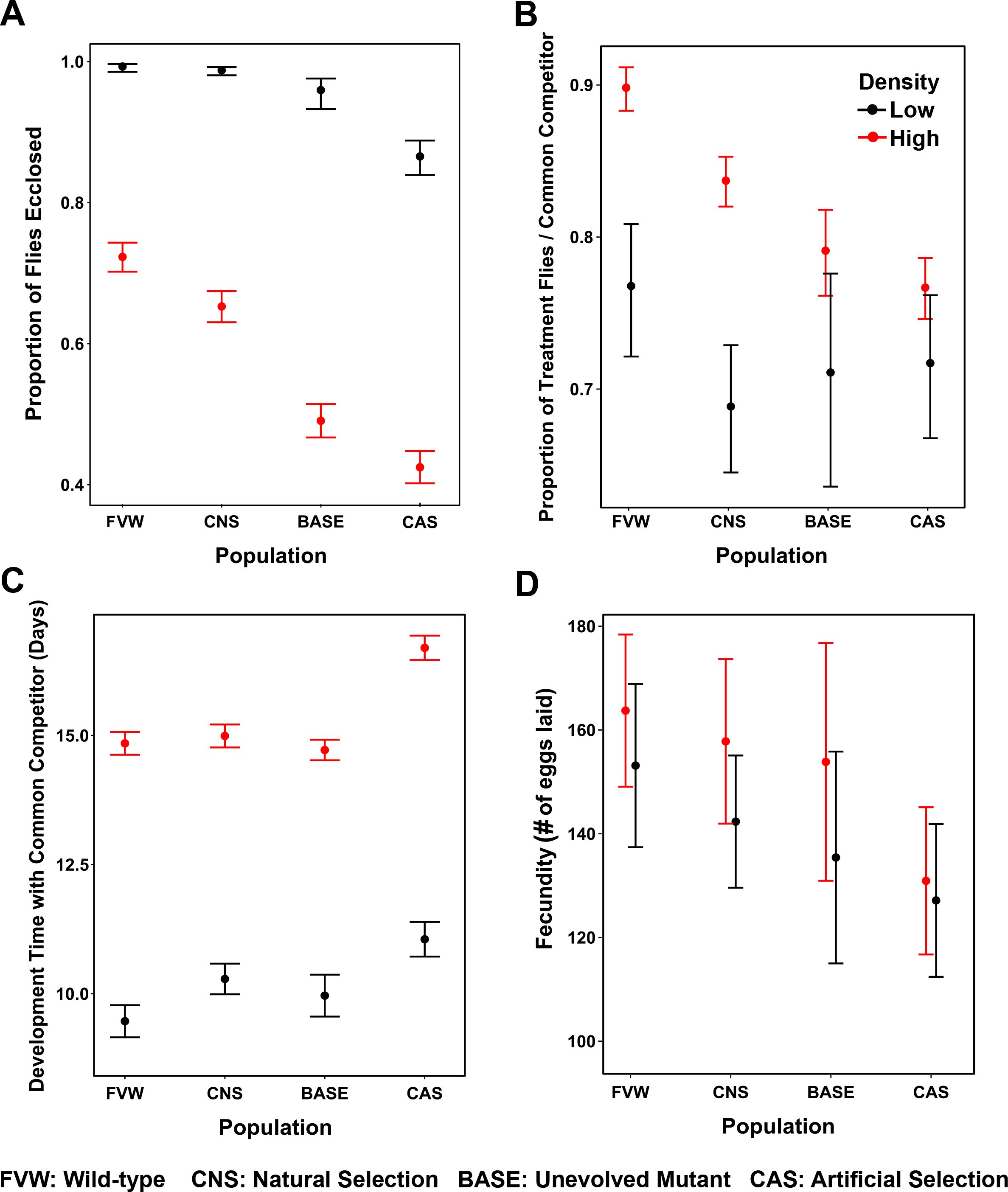
Trade-offs between morphological recovery and other fitness-associated (life-history) traits. (A) Egg to adult survivorship at low (50 eggs) and high (300 eggs) density, higher in CNS populations and lower in CAS populations as compared to mutant controls (B) Larval competitive ability in the presence of a common competitor at low (25+25 eggs) and high (150+150 eggs) density, higher especially at high density in CNS populations and lower in CAS populations as compared to mutant controls (C) Egg to adult development time in the presence of a common competitor at low and high density, higher especially at high density in CAS populations as compared to mutant controls. The data points for (A), (B) and (C) and error bars represent the respective mean phenotypes and 95% CI of 6 vials/ population/ replicate/ density (D) Fecundity when reared at low and high density, demonstrating no difference between the populations. The data points represent the mean fecundity and error bars represent the 95% CI, for 20 male-female pairs (one per vial) / population/ replicate/ density over 4 days of egg laying.

While the common competitor strain was generally weaker and survived poorly compared to most evolved populations, the natural selection lineages still demonstrated higher larval competitive ability at high density as compared to the BASE and the artificially selected CAS lineages (Fig. 4B and Fig. S6B). As observed for viability, the CAS lineages performed poorly in competition and with increased development time (Fig. 4C and Fig. S6C) compared with all other lineages at both high and low density. We performed an additional assay, where the experimentally evolved CNS and BASE populations were competed against FVW as the common competitor. The CNS populations were more competitive at both densities compared to BASE when competed against FVW (Fig. S7B). These results demonstrated that the CNS populations had compensated for some of the deleterious effects of the *vg*^*1*^ allele on competitive ability, while the artificially selected lineages showed reduced performance with respect to viability, development time and competitive ability. Interestingly, we saw no consistent pattern of evolved response for fecundity (Fig. 4D and Fig. S6D).

## Discussion

Compensatory evolution in microbial (4, 8–13) and other systems (5, 14–16) demonstrates the ubiquity of *de novo* compensatory mutations contributing to adaptation. But whether fitness recovery can occur via both direct and indirect effects on the phenotypes influenced by the deleterious mutation and how selection utilizes standing genetic variation for both phenotypic and fitness compensation is unclear. To address these, we fixed several mutations, in a large natural population of *D. melanogaster* and subjected them to independent regimes of artificial selection and natural selection. Focusing mainly on one of the mutations, *vg*^*1*^, during the course of selection we assayed and found evidence for rapid compensatory evolution in morphological, behavioural and life-history traits in these populations contingent upon the selection regime.

Wing morphology in *Drosophila* is a target of selection (26–29). We confirmed that *vg*^*1*^ allele had severe sexual selection dependent and independent deleterious effects on fitness that could be potentially compensated by selection (Fig. 2A and Fig. S2A). Furthermore when mate choice was present, sexual selection and natural selection acted potentially synergistically to accelerate the loss of the *vg*^*1*^ allele. Thus the naïve prediction might be that natural selection would compensate for the defects in wing morphology to ameliorate this effect.

The rapid recovery of almost completely wild type wing morphology in all replicates of CAS populations upon artificial selection (Fig. 1B) demonstrates the presence of compensatory alleles segregating in natural populations (22, 23). However the natural selection lineages did not compensate for the wing morphology defects caused by the introduced mutations and may even undergoing further, very gradual wing reduction (Fig. 1B). This suggests that the lineages only experiencing natural selection evolved along an alternative trajectory and potentially reduced the importance of wing-mediated fitness gain (25). While sexual selection was clearly still operating, other agents of selection on wing morphology (and performance) such as flight were likely highly relaxed. The degree to which the response in the naturally selected lineages would recapitulate that observed by artificial selection (including antagonistic pleiotropic effects on other fitness components) is an important area for further study.

One potential wing-independent trajectory of evolution involved modifying mating behaviour to by-pass wing-mediated sexual signalling. Copulation success and especially courtship rates were increased in the experimentally evolved CNS populations relative to other lineages (Fig. 3, Fig. S3, Fig. S4 and Fig. S5). This modification of courtship was largely due to increased male persistence (Fig. S3A and Fig. S4), demonstrating that the CNS populations had compensated for the lack of wing mediated sexual signalling via an increase in male courtship persistence in these populations. Our observations are similar to changes in *Teleogryllus oceanicus* in nature (17), where over 90% of males have a flatwing morphology to reduce parasitism. This flatwing renders male crickets unable to produce a mating call. Despite this, the frequency of the flatwing morphology appears to be maintained in the population due to evolution of modified mating behaviours (18, 19) compensating for loss of courtship song production. Interestingly the lineages artificially selected to compensate for the effects on wing morphology show higher rates of courtship and copulation than the experimentally evolved populations.

Given the mating costs associated with the *vg*^*1*^ allele, and segregating variation to compensate for defects, it may initially seem puzzling why experimentally evolved lineages do not compensate for wing morphology. This is resolved by examining the costs for other fitness components, where the experimentally evolved populations outperform the ancestral mutant, while the artificially selected lineages do worse. We observed that egg to adult viability and larval competitive ability in the experimentally evolved CNS populations increased compared to the BASE population and recovered to almost wild-type levels (Fig. 4A-B and Fig. S6A-B). This is consistent with other experiments testing adaptation to high larval density where egg to adult viability and larval competitive ability evolved (34, 35). In contrast, the lineages that recovered “wild type” wing morphology under artificial selection (CAS) had reduced egg to adult survivorship, larval competitive ability and increased development time (Fig. 4A-C, Fig. S6A-C), compared even to the BASE and the population size matched but unselected NASC lineages. This suggests that the alleles (or linked alleles) that provide phenotypic compensation of the wing phenotype might have negative pleiotropic consequences reflected in other fitness components.

We performed similar selection with mutations in *rhomboid* (*rho*^*ve-1*^) and *net* (*net*^*1*^) genes. Mutations with similar phenotypic consequences are present in low frequencies in natural populations (33) and previous selection experiments with these have produced rapid phenotypic response by artificial selection (24, 36–38). We confirm these results, and show that segregating compensatory modifiers can mediate rapid suppression of the mutant phenotypes in these populations under artificial selection (Fig. 1A). Like our observations with the more severe *vg*^*1*^ allele, the experimentally evolved CNS lineages fixed for *rho*^*ve-1*^ and *net*^*1*^ mutations do not recover wild type like wing morphology. While *vg*^*1*^ is a strong perturbation with severe fitness consequences, these alleles have relatively weak phenotypic effects with respect to wing morphology and likely have smaller effects on fitness. Thus it is clear that natural populations are segregating variation that can compensate for a variety of deleterious mutational effects.

In conclusion, the artificial selection lineages rapidly recovered from the focal defect in the wing phenotype consequently recovering mating ability but have lower survivorship and larval competitive ability. The natural selection lineages seem to have taken an alternative evolutionary trajectory by modifying courtship behaviour to compensate for the loss of wing-mediated sexual signalling as well as increasing survivorship and larval competitive ability without recovery of the wing phenotype. Thus our study demonstrates fitness recovery can be mediated via direct or indirect compensation of multiple phenotypic aspects perturbed by a mutation and that rapid and consistently repeatable compensatory evolution can occur from standing genetic variation across multiple mutations (with severe or weak effects). Our results provide an interesting perspective on “Dollo’s law” on the irreversibility of evolution of trait loss (39–43). At least in the short term, the evolutionary loss of a trait may remain easily reversible (44–47) with the acquisition of compensatory mutations, including those concurrently segregating in natural populations. However, associated with trait loss (and any loss with respect to specific fitness components) may be new evolutionary opportunities that fundamentally alter the evolutionary trajectory of the populations in question (11, 25). This may mean that after a relatively small number of generations, the question is not whether there is a genetic constraint on the re-acquisition of the trait in question, but whether fitness surface has been sufficiently altered that selection would not favour trait reacquisition.

## Acknowledgements

Reuven Dukas, John Pool and Arnar Palsson provided valuable feedback on previous drafts. This work was supported by the BEACON center for the study of evolution in action (National Science Foundation) under Cooperative Agreement No. DBI-0939454 and MCB-0222344, NSERC Discovery (Canada) and funds from Michigan State University to ID.

---

**Supplementary Fig. 1.**
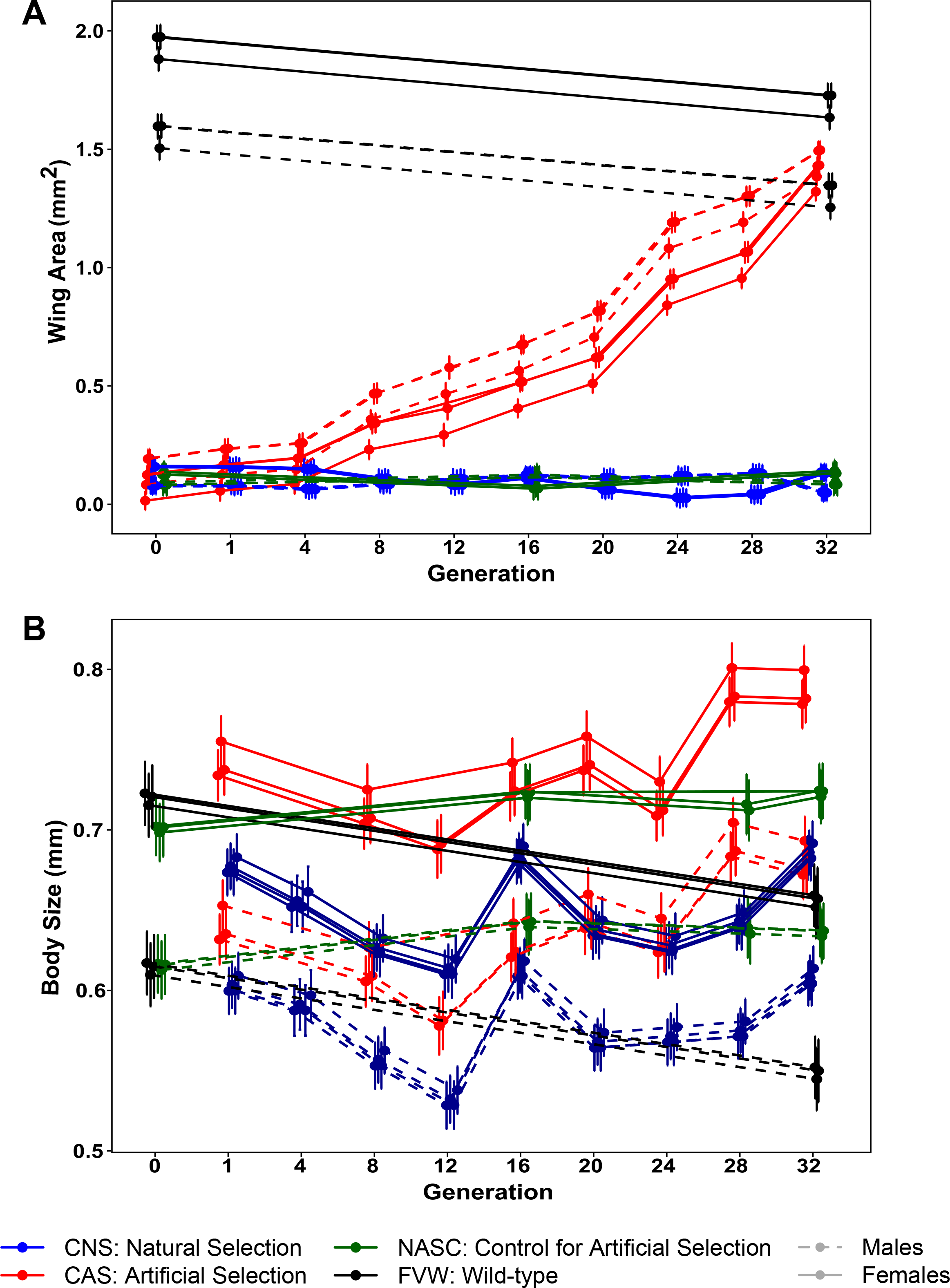

**Supplementary Fig. 2.**
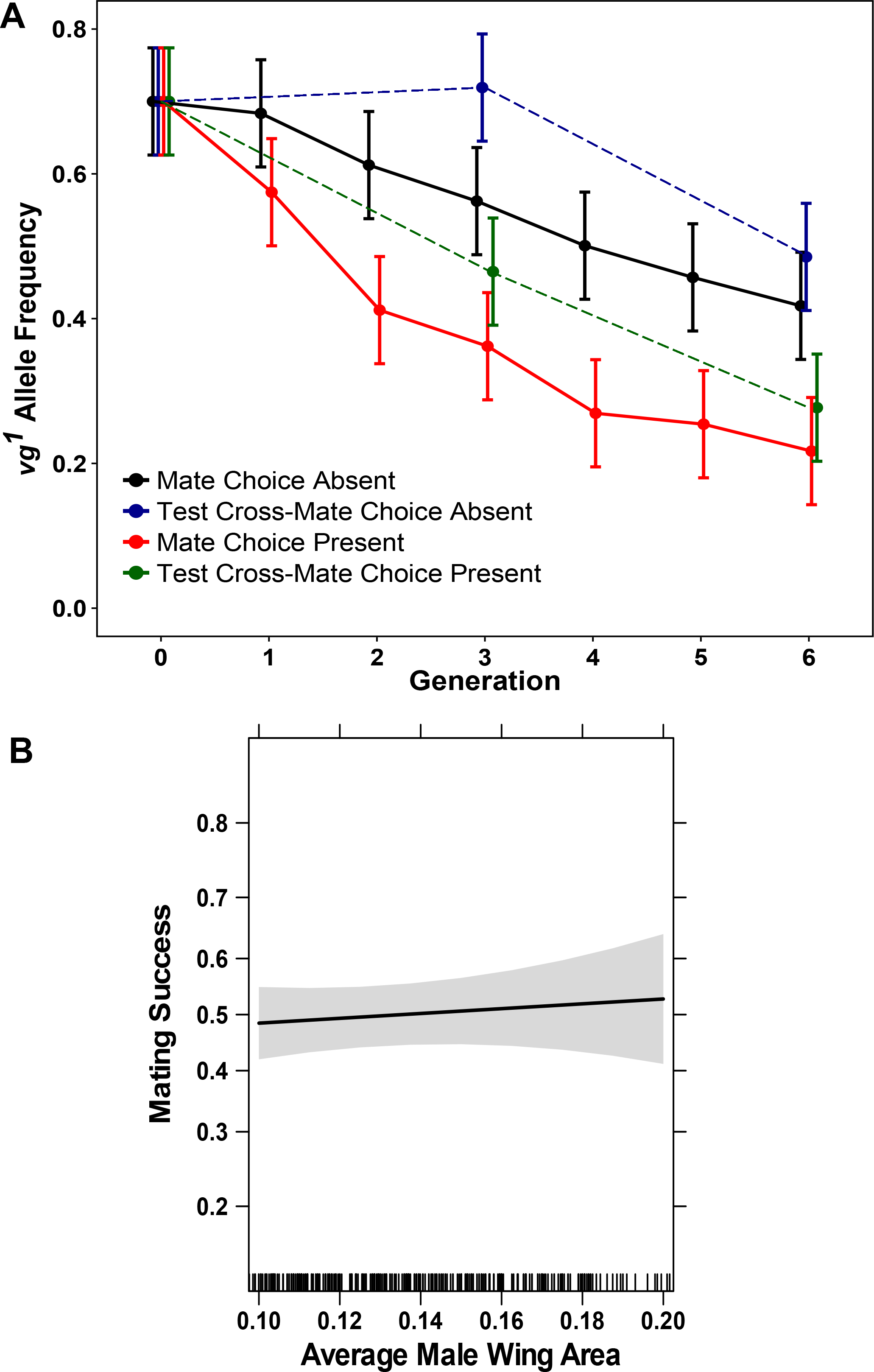

**Supplementary Fig. 3.**
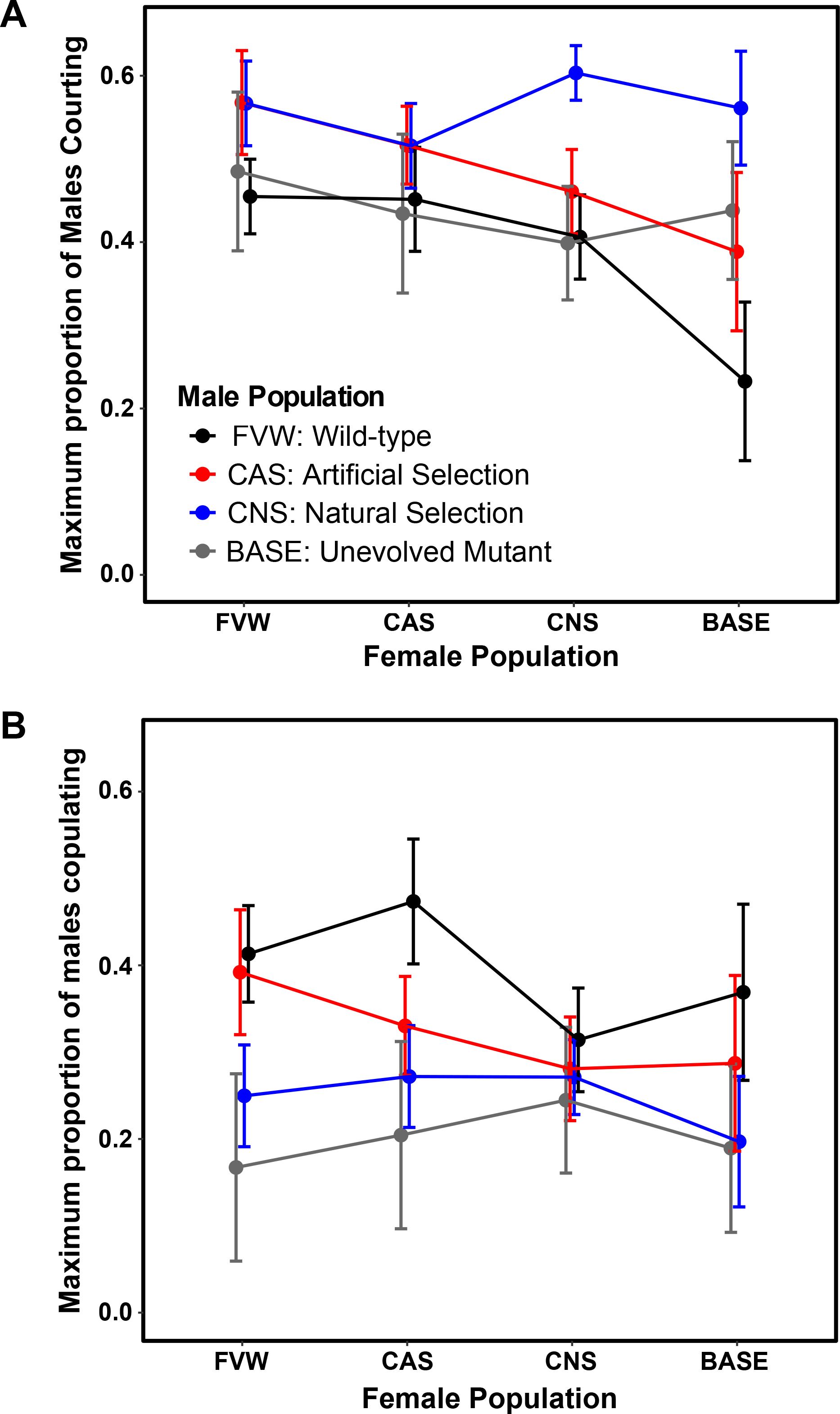

**Supplementary Fig. 4.**
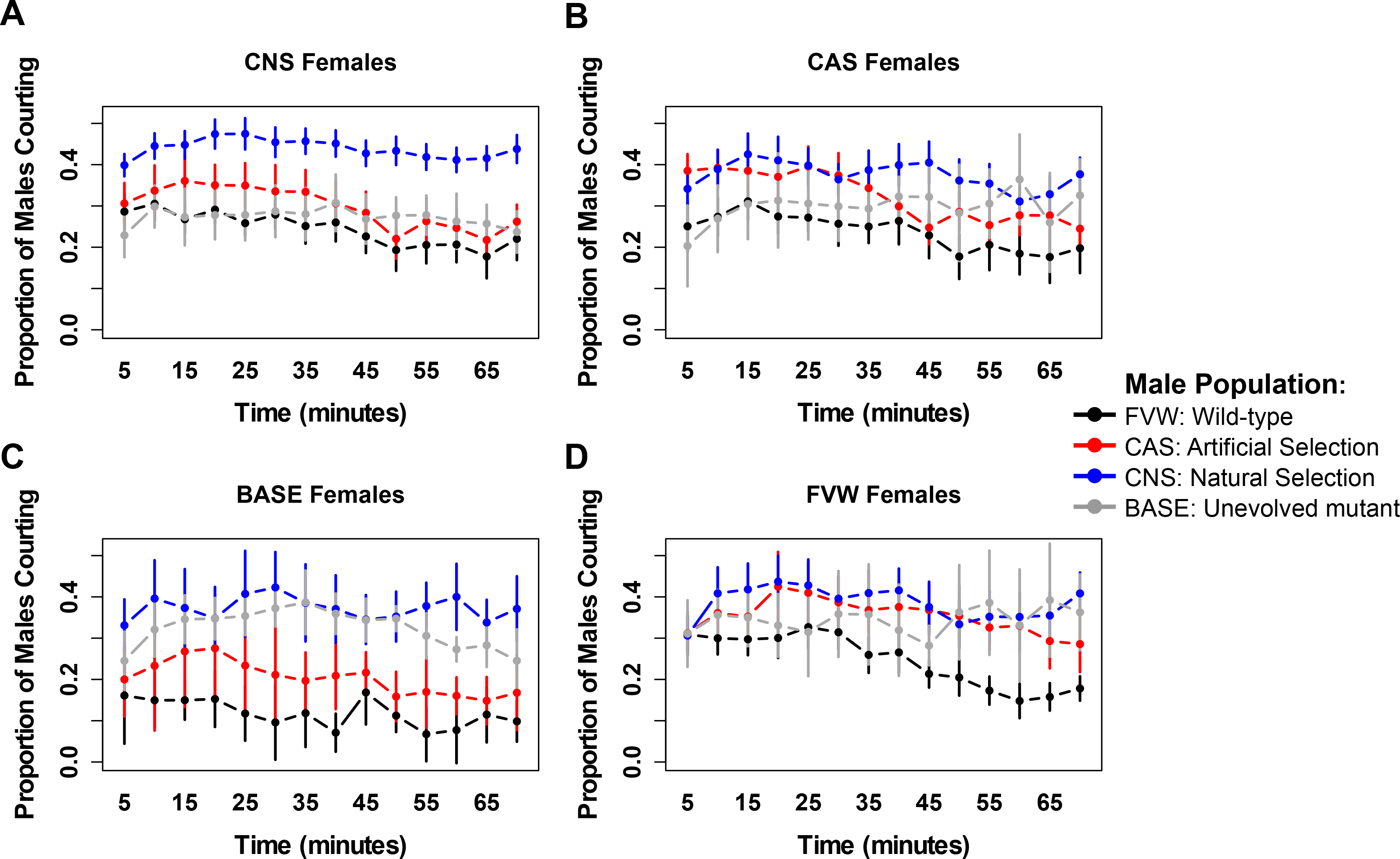

**Supplementary Fig. 5.**
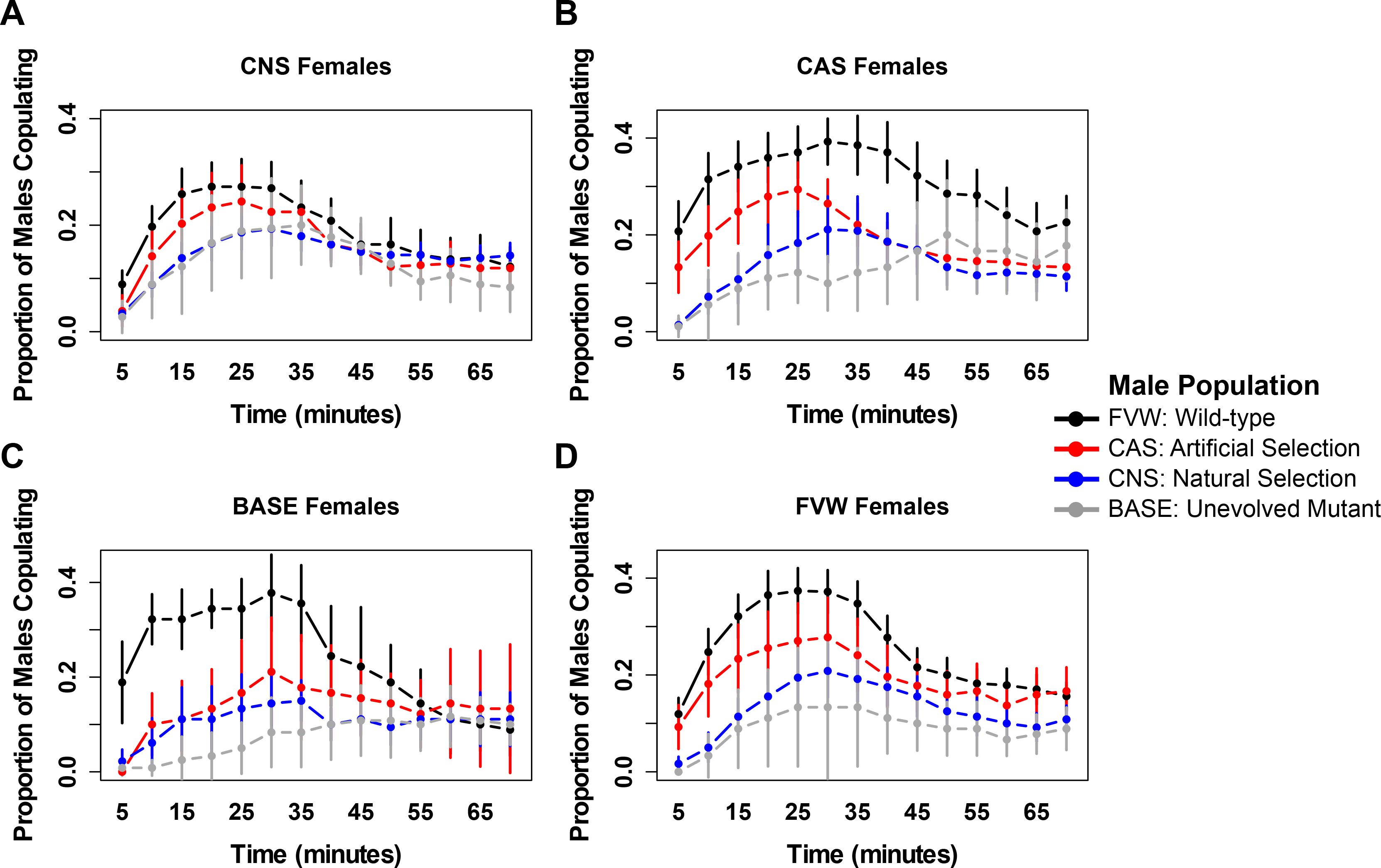

**Supplementary Fig. 6.**
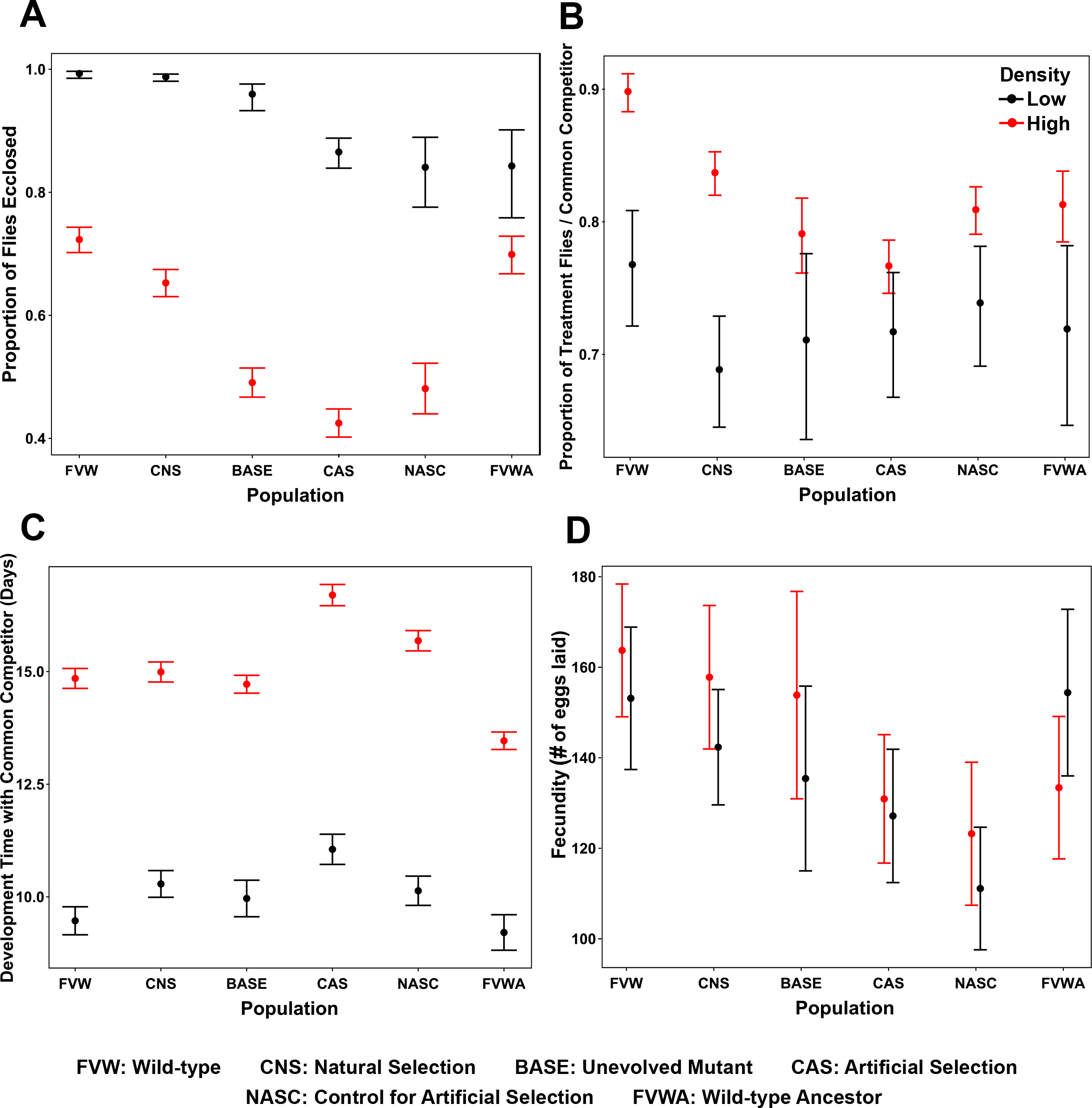

**Supplementary Fig. 7.**
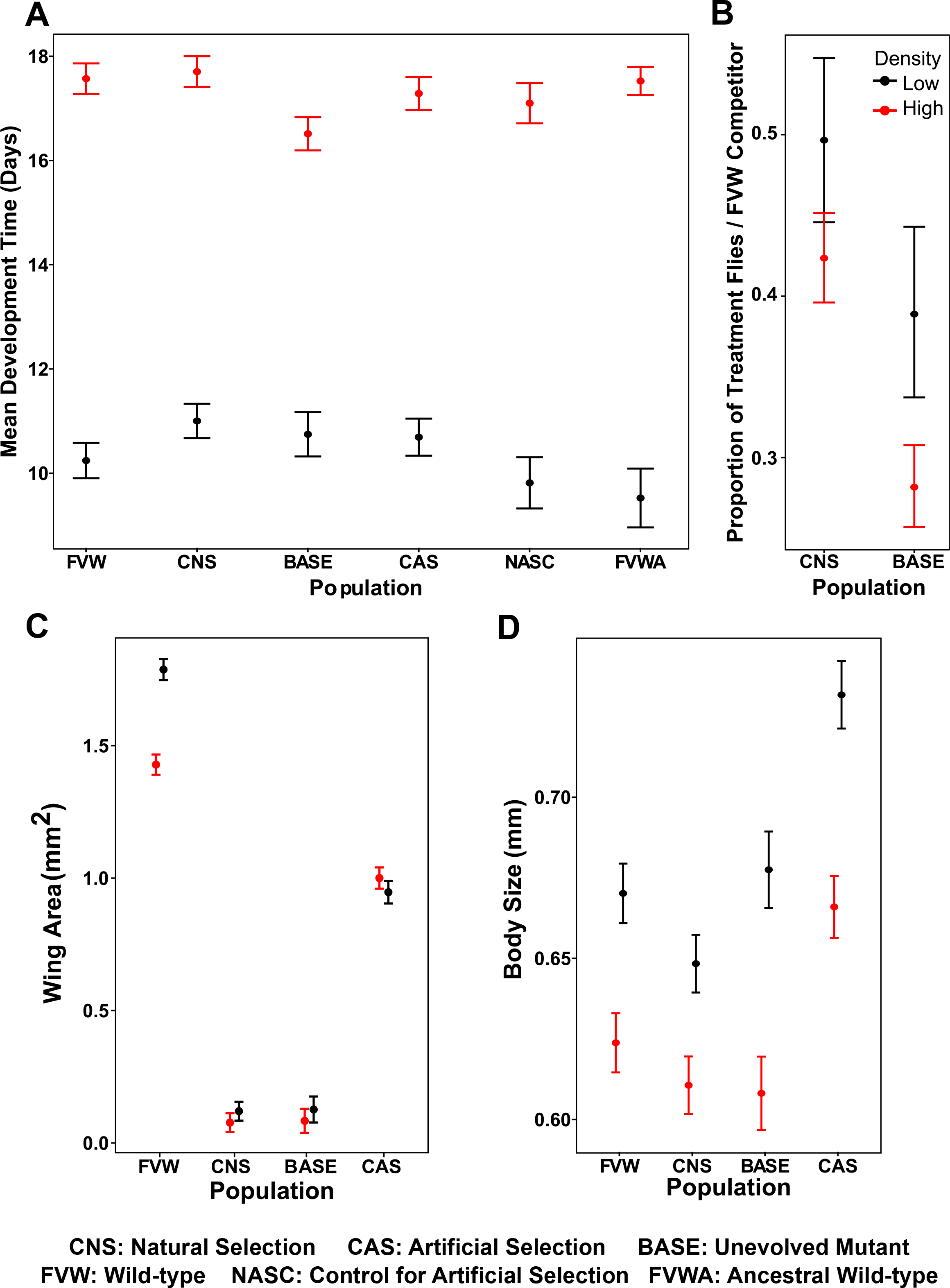

